# Dynamics of the Ghrelin/Growth Hormone Secretagogue Receptor System in the Human Heart Before and After Cardiac Transplantation

**DOI:** 10.1101/362525

**Authors:** Rebecca Sullivan, Varinder K Randhawa, Anne Stokes, Derek Wu, Tyler Lalonde, Bob Kiaii, Leonard Luyt, Gerald Wisenberg, Savita Dhanvantari

## Abstract

**Background:** Currently, the early pre-clinical detection of left ventricular (LV) dysfunction is difficult as biomarkers are not specific for the cardiomyopathic process. The underlying molecular mechanisms leading to heart failure remain elusive, highlighting the need for identification of cardiac-specific markers. The growth hormone secretagogue receptor (GHSR) and its ligand ghrelin are present in cardiac tissue and are known to contribute to myocardial energetics. Here, we examined tissue ghrelin-GHSR levels as specific markers of cardiac dysfunction in patients who underwent cardiac transplantation.

**Methods and Results:** Samples of cardiac tissue were obtained from 10 cardiac transplant patients at the time of organ harvesting, and during serial post-transplant biopsies. Quantitative fluorescence microscopy using a novel fluorescent ghrelin analog was used to measure levels of GHSR, and immunofluorescence was used to measure levels of ghrelin, b-type natriuretic peptide (BNP) and tissue markers of cardiomyocyte contractility and growth. GHSR and ghrelin expression levels were highly variable in the explanted heart, less in the grafted heart biopsies. GHSR and ghrelin were strongly positively correlated, and both markers were negatively correlated with LV ejection fraction. Ghrelin had stronger positive correlations than BNP with the signaling markers for contractility and growth.

**Conclusions:** These data suggest that GHSR-ghrelin have potential use as an integrated marker of cardiac dysfunction. Interestingly, tissue ghrelin appeared to be a more sensitive indicator than BNP to the biochemical processes that are characteristic of heart failure. This work allows for further use of ghrelin-GHSR to interrogate cardiac-specific biochemical mechanisms in pre-clinical stages of HF.

**Precis:** This study shows the relationships between GHSR, ghrelin, and signaling molecules with relation to heart function in human heart failure with tissue from diseased heart and healthy heart biopsies.

## 1. Introduction

The peptide hormone ghrelin is well-known as a potent orexigenic hormone. It stimulates food intake by activating hypothalamic neurons that regulate normal feeding behaviour^1^. It is the natural ligand of the growth hormone secretagogue receptor 1a (GHSR1a), a seven transmembrane, G protein-coupled receptor, which, in addition to the hypothalamus, is expressed in other brain regions as well as several endocrine organs, such as the anterior pituitary, pancreatic islets, the intestine, thyroid, and adipose tissue. In addition, ghrelin and GHSR are both expressed in cardiomyocytes where they function through an axis that is independent of their role in regulating energy expenditure^2^. Activation of GHSR in cardiomyocytes promotes excitation-contraction coupling by increasing Ca^2+^ flux through both voltage-dependent Ca^2+^ channels^3^ and the sarcoplasmic reticulum Ca^2+^-ATPase pump (SERCA2a)^4–6^, and promotes cardiomyocyte growth and survival through extracellular signaling related kinase1/2 (ERK1/2)^4,5^, and phosphatidylinositol-3-kinase / Akt (PI3K/AKT)^5,7^. We^8^, and others^3^, have recently shown that levels of GHSR are decreased in rodent models of diabetic cardiomyopathy, thus suggesting that the myocardial ghrelin/GHSR system could be a biomarker for cardiomyopathies.

Similarly, levels of ghrelin and GHSR are dramatically altered throughout the heart in patients with severe heart failure ^9^. The clinical syndrome of heart failure (HF) is most commonly associated with significant impairment of left ventricular (LV) contractility, leading to elevated intra-cardiac diastolic pressures and extravasation of fluid into the lung parenchyma and other tissues. The early detection and treatment of HF are limited by two issues: a) the specific series of molecular mechanisms leading to impaired contractility remain elusive in patients with idiopathic cardiomyopathies, and b) the responses to guideline-directed medical therapies remain highly variable, such that many patients continue to deteriorate, leading to either the need for cardiac transplantation or ultimately death. Clinically, there is a critical need to prospectively identify groups of patients who will ultimately be at higher risk, particularly in the early stages of left ventricular dysfunction, when the clinical status and ventricular function are not by themselves consistent reliable predictors of disease progression and clinical outcomes. Circulating biomarkers, such as natriuretic peptide type-B (BNP), particularly the N-terminal form (NT-proBNP), and troponins T and I^10–12^, provide some prediction of the progression of HF, by indicating changes within the cardiomyocyte that lead to stress, injury and apoptosis. However, they are produced whenever heart tissue is damaged by any direct or indirect injury to the myocardium, and there may be discordance between tissue and circulating levels of these biomarkers. Therefore, there is a need to identify myocardial-specific biomarkers that reflect the cellular and molecular processes that underlie the progression of HF.

In this study, we evaluated the role of tissue ghrelin-GHSR levels as a specific marker of cardiac dysfunction in a cohort of patients who underwent cardiac transplantation. We examined samples from the diseased explanted heart and biopsies from the same individual’s healthy donor heart followed through to one-year post-transplantation. In addition, we aimed to determine the relationship between ghrelin-GHSR and BNP to biochemical signaling molecules in cardiac dysfunction. Given the importance of intracellular Ca^2+^ homeostasis in atrial and ventricular contractility and the role of phospho-ERK 1/2 in cardiomyocyte growth, we hypothesized that changes in the ghrelin-GHSR axis in myocardial tissue could potentially reflect derangements in cardiomyocyte contractility and initiation of cardiac hypertrophic reprogramming that characterize the progression of heart failure.

## 2. Materials and methods

### 2.1 Patient Cohort

Tissue samples were harvested from 10 patients who underwent cardiac transplantation at the London Health Sciences Center (LHSC) between 2011 and 2013. The protocol for sample dissections was approved by Western University’s Health Sciences Research Ethics Board. Samples, roughly 0.5cm to 1.5cm in length, were collected from the right atrium (RA) and left ventricle (LV) of the explanted/diseased heart (DH) from each cardiac transplant patient. Endomyocardial biopsies, roughly 0.1 to 0.3cm in length, from the right ventricle (RV) of the newly grafted heart were also taken at various time-points post-transplantation (PTx), generally weekly for the first 4 weeks, monthly for months 2-6 and then at 1 year PTx. Patient demographics, cardiac function (LVEF), and medications pre- and post-transplantation are shown in Tables 1 and 2, respectively. All patient samples and patient data were kept anonymous and all marker analyses was done prior to receiving clinical data.

**Table 1.** Information on antibodies used.

| Antigen | Catalog # | Dilution | Host | RRID# |
| --- | --- | --- | --- | --- |
| Ghrelin | sc-10359 | 1:100 | Goat | AB_2111733 |
| Serca2A | ab3625 | 1:300 | Rabbit | AB_303961 |
| pERK1/2 | sc-377400 | 1:250 | Mouse |  |
| BNP | ab19645 | 1:1000 | Rabbit | AB_445037 |
| Collagen I | ab34710 | 1:500 | Rabbit | AB_731684 |
| DAPI | 62247 | 1:1000 | - | AB_2629482 |
| Alexafluor 488 | A11055 | 1:500 | Donkey anti-goat | AB_2534102 |
| Alexafluor 594 | A21207 | 1:500 | Donkey anti-rabbit | AB_141637 |
| Alexafluor 594 | A21203 | 1:500 | Donkey anti-mouse | AB_141633 |
| Alexafluor 488 | A21206 | 1:500 | Donkey anti-rabbit | AB_141708 |

**Table 2.**
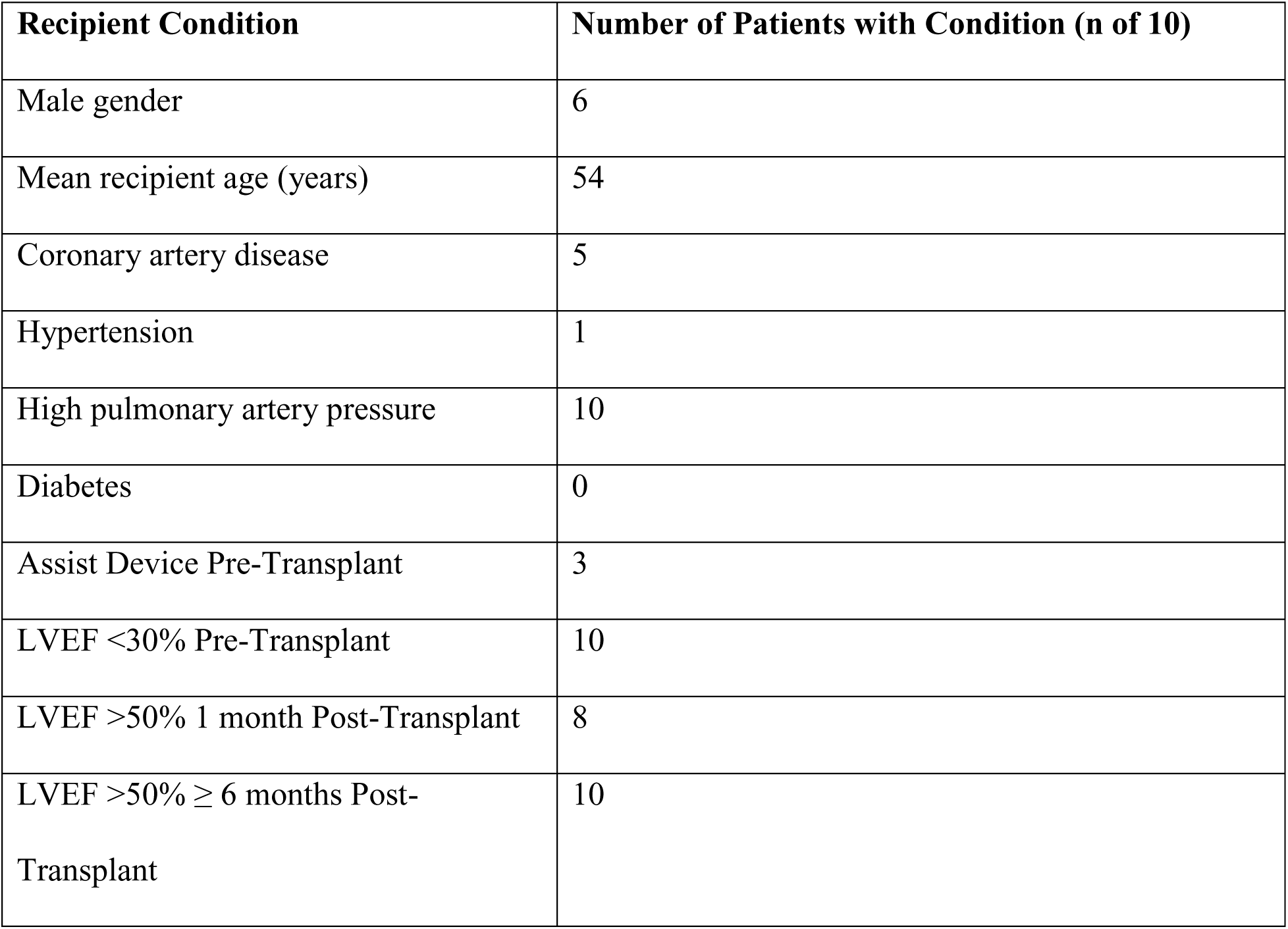
Cardiac Transplant Recipient Patient Demographics

| **Recipient Condition** | **Number of Patients with Condition (n of 10)** |
| --- | --- |
| Male gender | 6 |
| Mean recipient age (years) | 54 |
| Coronary artery disease | 5 |
| Hypertension | 1 |
| High pulmonary artery pressure | 10 |
| Diabetes | 0 |
| Assist Device Pre-Transplant | 3 |
| LVEF <30% Pre-Transplant | 10 |
| LVEF >50% 1 month Post-Transplant | 8 |
| LVEF >50% ≥ 6 months Post-Transplant | 10 |

### 2.2 Immunofluorescence microscopy

Samples from both the explanted (diseased) and grafted hearts were frozen and embedded in optimal cutting temperature compound (OCT), and subsequently sectioned at 7 μm thickness, as previously described ^4,13^. Immunohistochemistry using fluorescent antibodies was conducted as previously described ^4,13^. In brief, tissue sections were incubated with primary polyclonal or monoclonal antibodies (Table 1) for 1h at room temperature in a humidified chamber. These antibodies were used to identify ghrelin (1:100), BNP (1:1000), pERK1/2 (1:250); SERCA2a (1:300), and collagen I (1:500). Samples were rinsed twice in phosphate buffered solution (PBS) and incubated for 2h at room temperature with secondary antibodies (1:500) (Table 1). We have previously used the far-red ghrelin analog probe (Ghrelin(1-18, Lys^18^(Cy5)) to quantify GHSR *in situ*^4^. This analog binds with high specificity to GHSR in mouse cardiac tissue samples^6^. Therefore, we used Ghrelin(1-18, Lys^18^(Cy5) to quantify the expression of GHSR. Following incubation with secondary antibodies, this probe was added to tissue sections for 30 min and used for GHSR detection. Sections were washed with PBS, incubated 8 min with DAPI nuclear stain (1:1000), and mounted with ProLong Gold antifade (Life Technologies) to prevent the tissues from photobleaching. Images were captured with a Nikon Eclipse TE2000-S fluorescent microscope. Five random fields of view were acquired for each of 4 tissue sections at 20X magnification (Nikon NIS Elements v. BR 4.50.00).

### 2.3 Fibrosis imaging

In order to assess fibrosis, heart tissue sections were stained with Masson’s Trichrome stain by the core Pathology laboratory at LHSC. Sections were acquired using bright field microscopy at 10X, 20X or 40X magnifications with a Zeiss Axioskop EL-Einsatz microscope and Northern Eclipse software. Fluorescence microscopy was also used to acquire collagen I images as described above.

### 2.4 Data Analysis

Images of GHSR, ghrelin, and biochemical signaling molecules were analyzed with FIJI v. 1.49v, a distribution of ImageJ software (National Institutes of Health, Bethesda). Fluorescence of each section was quantified using a custom FIJI script that integrates raw density images that represent protein expression levels, as previously reported by us^4,14^. Briefly, thresholding was conducted for each image to determine the fluorescence intensity (positive pixel count above threshold minus the background). Fibrosis was analyzed using an online script which quantified the percentage of fibrotic tissue in each sample by distinguishing fibrotic tissue from non-fibrotic tissue^15^. Statistical analyses were performed using GraphPad Prism version 7.02 or IBM SPSS statistics 25, as follows: unpaired student *t* test, a one-way ANOVA with analysis of variance using Tukey *post-hoc* test to compare differences between diseased hearts and biopsies of the grafted hearts; Pearson correlation and logistic linear regression for correlations between markers: and Spearman bivariate correlation for relationships between LVEF and the following markers: GHSR, ghrelin, BNP, pERK1/2 and SERCA2a, all with significance set at p<0.05.

## 3. Results

### 3.1 Cardiac transplant patient cohort

Sixty percent of the patients who underwent cardiac transplantation were male, with an overall mean age of 54 years (Table 2). Fifty percent had significant coronary artery disease pre-transplant, and none had diabetes. All patients had elevated pulmonary artery pressures pre-transplant, along with severely reduced left ventricular ejection fraction (LVEF <30%) by echocardiographic assessment. Serial echocardiographic assessment of LV function following cardiac transplant showed LV recovery to an LVEF > 50% in 80% of patients by 1 month PTx. By 6 months PTx, all patients had normal LV function (Table 2). Although most patients were receiving HF medications, i.e., angiotensin converting enzyme (ACE) inhibitors/beta blockers/diuretics, prior to transplantation, only a small number continued to receive these medications following surgery. Table 3 contains a complete listing of medications pre- and post-Tx.

**Table 3.** Patient Medications Pre- and Post-Cardiac Transplant.

| **Medications** | **Pre-transplant**<br>**(n of 10)** | **Post-transplant**<br>**1 month (n of**<br>**10)** | **Post-transplant**<br>**6 months (n of**<br>**10)** | **Post-transplant**<br>**1 year (n of 10)** |
| --- | --- | --- | --- | --- |
| ACE Inhibitor * | 8 | 2 | 2 | 8 |
| Anti-arrhythmic | 5 | 2 | 0 | 0 |
| Angiotensin<br>receptor blocker<br>(ARB) | 0 | 0 | 0 | 0 |
| Anti-platelet | 5 | 7 | 7 | 8 |
| Coumadin | 7 | 1 | 0 | 0 |
| Beta blocker | 10 | 1 | 0 | 0 |
| Calcium channel<br>blocker | 10 | 2 | 4 | 3 |
| Digoxin | 8 | 0 | 0 | 0 |
| Diuretics | 10 | 7 | 2 | 2 |
| Statins | 5 | 5 | 6 | 6 |
| Nitrates | 0 | 0 | 0 | 0 |
| Nitroglycerin | 0 | 0 | 0 | 0 |
| Anti-diabetics | 0 | 0 | 0 | 0 |
| Inotropic<br>support | 1 | 0 | 0 | 0 |
| (Milrinone /<br>Dobutamine) |  |  |  |  |
| Tacrolimus | 0 | 9 | 10 | 10 |
| Mycophenolate<br>mofetil | 0 | 9 | 9 | 9 |
| Prednisone | 0 | 7 | 9 | 6 |

### 3.2 GHSR and ghrelin expression in cardiomyocytes

In any given cardiac transplant patient, the expression of GHSR and ghrelin in both LV and RA appeared to be elevated in the explanted heart in comparison to their expression PTx in the grafted heart tissue biopsies over time (Figure 1 A-C). By logistic regression analysis of all patient specimens, both pre-and post-transplant, the level of GHSR expression demonstrated a strong and highly significant positive correlation with the level of ghrelin expression (r = 0.7817, *p*<0.0001). To examine associations of ghrelin and GHSR with cardiac dysfunction, data were divided into 2 groups of LVEF < 30% (pre-transplant) and LVEF > 50% (post-transplant), as there were no midrange values of LVEF. In the pre-transplant hearts with LVEF < 30%, expression of GHSR and ghrelin clustered towards the higher end of the regression line, while in grafted heart biopsies at 1 and 6 months (LVEF > 50%), expression of GHSR and ghrelin clustered towards the lower end of the regression line (Figure 1 D). To more closely examine correlations between ghrelin/GHSR and cardiac dysfunction, we used a Spearman bivariate correlation test, and showed significant negative correlations between GHSR and LVEF (p=0.018) and ghrelin and LVEF (p=0.004) (Figure 1 E).

**Figure 1.**
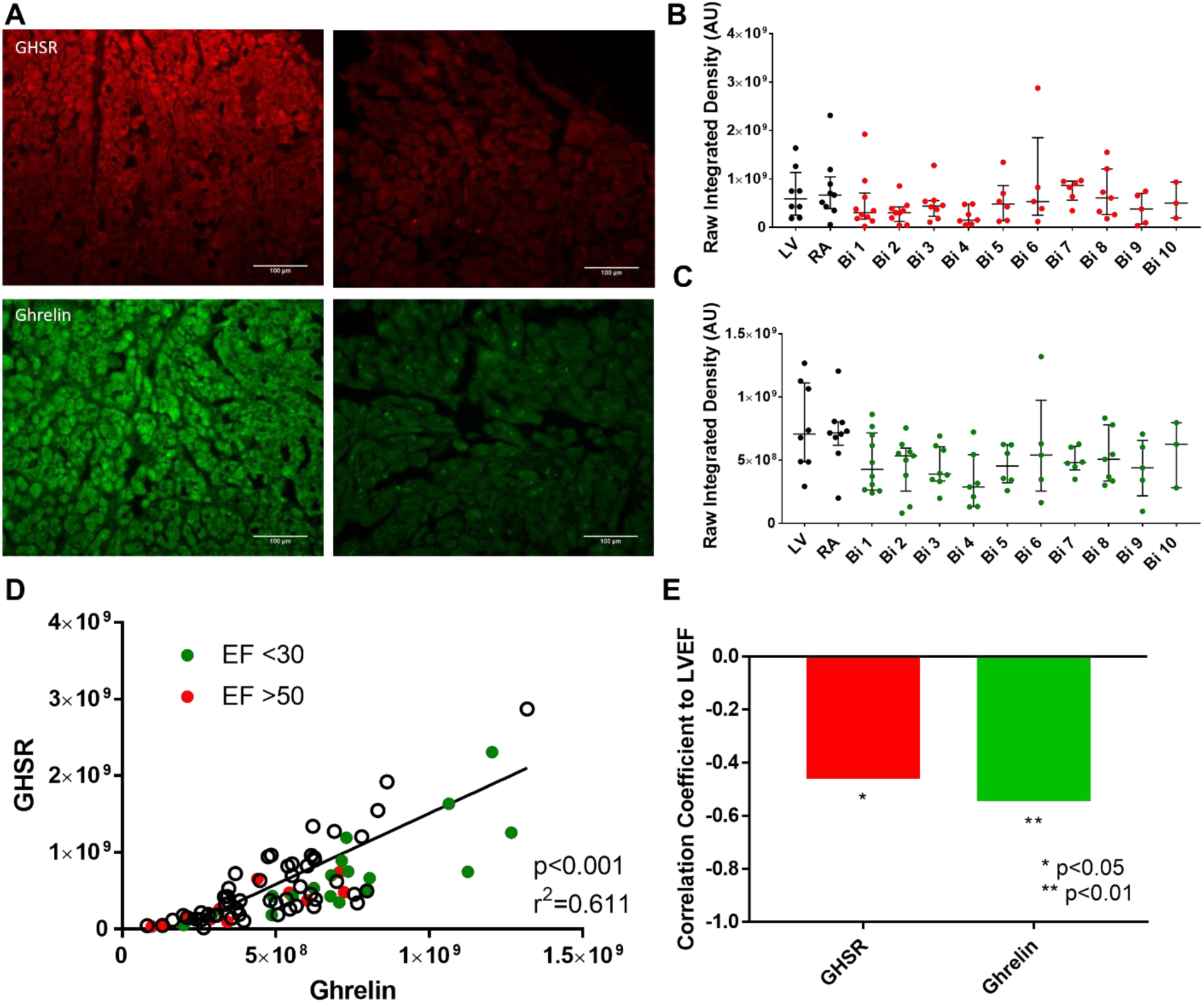
Ghrelin and GHSR expression in patients pre- and post- cardiac transplant. A) Representative fluorescent images of GHSR (red) and ghrelin (green) are shown in explanted heart tissue (left) and grafted heart biopsies (right) taken from the same patient. B and C) Quantified fluorescence intensity of GHSR (red) and ghrelin (green) is shown from explanted heart tissue (LV and RA, black dots) and grafted heart biopsies (Bi 1-10, coloured dots). D) Positive correlation of GHSR and ghrelin expression is shown in the entire cardiac transplant cohort with EF <30 in green and EF>50 in red. Each dot represents one transplant patient sample. LV: Left Ventricle; RA: Right atrium; Bi 1 – Bi 10: Biopsy 1 – Biopsy 10; EF: ejection fraction.

### 3.3 Metabolic markers in cardiac tissue

The expression of metabolic markers in cardiac tissue that are associated with cardiac dysfunction were measured by immunofluorescence microscopy. We measured the following markers: a) SERCA2a, as an index of cardiomyocyte contractility, b) pERK1/2, as a marker of cardiomyocyte growth/hypertrophy, and c) BNP, as a validated clinical tool used to determine the presence and degree of HF. Representative images of SERCA2a, pERK1/2 and BNP expression are shown in Figures 2A and 3A, respectively. The expression of SERCA2a was significantly elevated in cardiac tissue from the RA of the diseased heart (Figure 2 B) when compared to its expression in endomyocardial biopsies taken at weeks 1, 2, and 3 PTx by 3.2, 2.4, and 2.7 fold, respectively. However, there was no significant difference in SERCA2a expression in the RA after the first month PTx. There were also no significant differences in the expression of SERCA2a in the LV tissue of the diseased heart when compared to the PTx endomyocardial biopsies. pERK1/2 expression was significantly increased in the cardiac tissue of the diseased heart taken from the RA by 1.8-3.1-fold and LV by 1.7-2.9-fold when compared to the grafted heart tissue biopsies taken at any time-point PTx (Figure 2 C). The degree of BNP expression (Figure 3B) showed similar trends as for GHSR and ghrelin. Logistic regression analyses were performed to determine the association between SERCA2a, pERK1/2, GHSR and ghrelin levels (Figure 2). Highly significant and strong positive correlations were found between pERK1/2 and SERCA2a (r = 0.6867, *p*<0.0001), pERK1/2 and ghrelin (r = 0.5719, *p*<0.0001), and SERCA2a and ghrelin (r = 11 0.7171, *p*<0.0001). By contrast, there was a much weaker correlation between SERCA2a and GHSR (r = 0.2320, *p*=0.0483), and there was no correlation between pERK1/2 with GHSR. Logistic regression analyses were performed between all metabolic markers and BNP to determine any possible relationships (Figure 3 C-F). There was a positive correlation between ghrelin and BNP (r = 0.6782, *p*<0.0001), and SERCA2a and BNP (r = 0.4838, *p*<0.0001). However, there were weak correlations between GHSR and BNP (r = 0.2423, *p*=0.0282), and pERK1/2 and BNP (r = 0.2745, *p*=0.0196). Spearman bivariate correlations (correlation coefficients, CorC) were calculated as above and indicated highly significant negative associations between LVEF and SERCA2a (*p*<0.001, CorC = −0.63), and pERK (*p* < 0.001, CorC = −0.814), and between LVEF and BNP (*p* < 0.001, CorC = −0.773).

**Figure 2.**
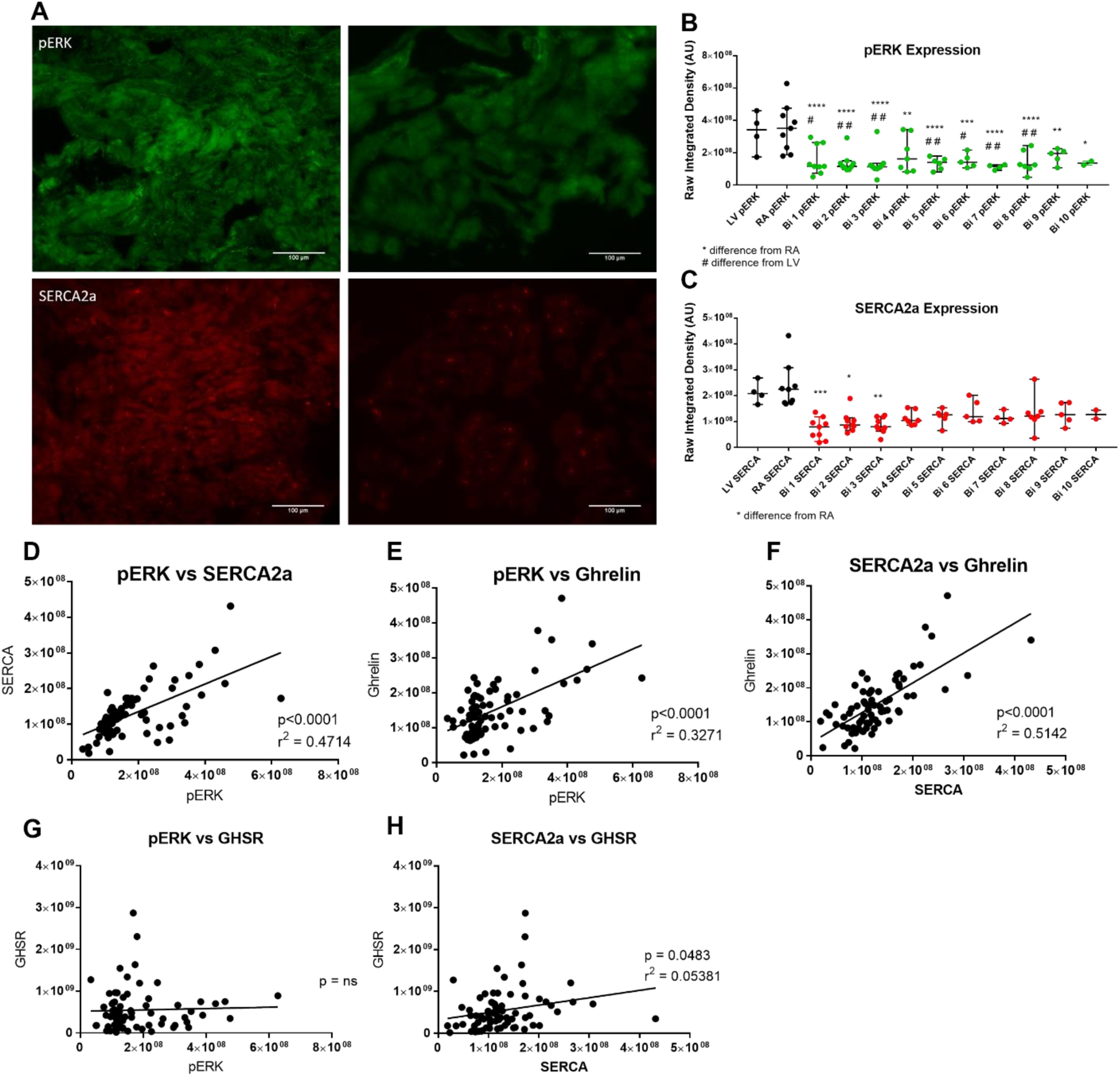
Cardiac metabolic markers in patients pre- and post-cardiac transplant in entire transplant patient cohort. A) Representative fluorescent images of pERK1/2 (green) and SERCA2a (red) are shown from explanted heart tissue (left) and grafted heart biopsies (right) samples taken from the same patient. B and C) Quantified fluorescence intensity of pERK1/2 (green) and SERCA2a (red) are shown for explanted heart tissue (LV and RA, black dots) and grafted heart biopsies (Bi 1-10, coloured dots). D-H) Positive correlation is seen in the entire cardiac transplant cohort for pERK1/2 vs SERCA2a, pERK1/2 vs ghrelin, SERCA2a vs ghrelin, pERK1/2 vs GHSR, and SERCA2a vs GHSR. Each dot represents one transplant patient sample. LV: Left Ventricle; RA: Right atrium; Bi 1 – Bi 10: Biopsy 1 – Biopsy 10. \**p*<0.05 from RA; ** *p*<0.01 from RA; *** *p*<0.001 from RA; **** *p*<0.0001 from RA; # *p*<0.05 from LV; ## *p*<0.01 from LV.

**Figure 3.**
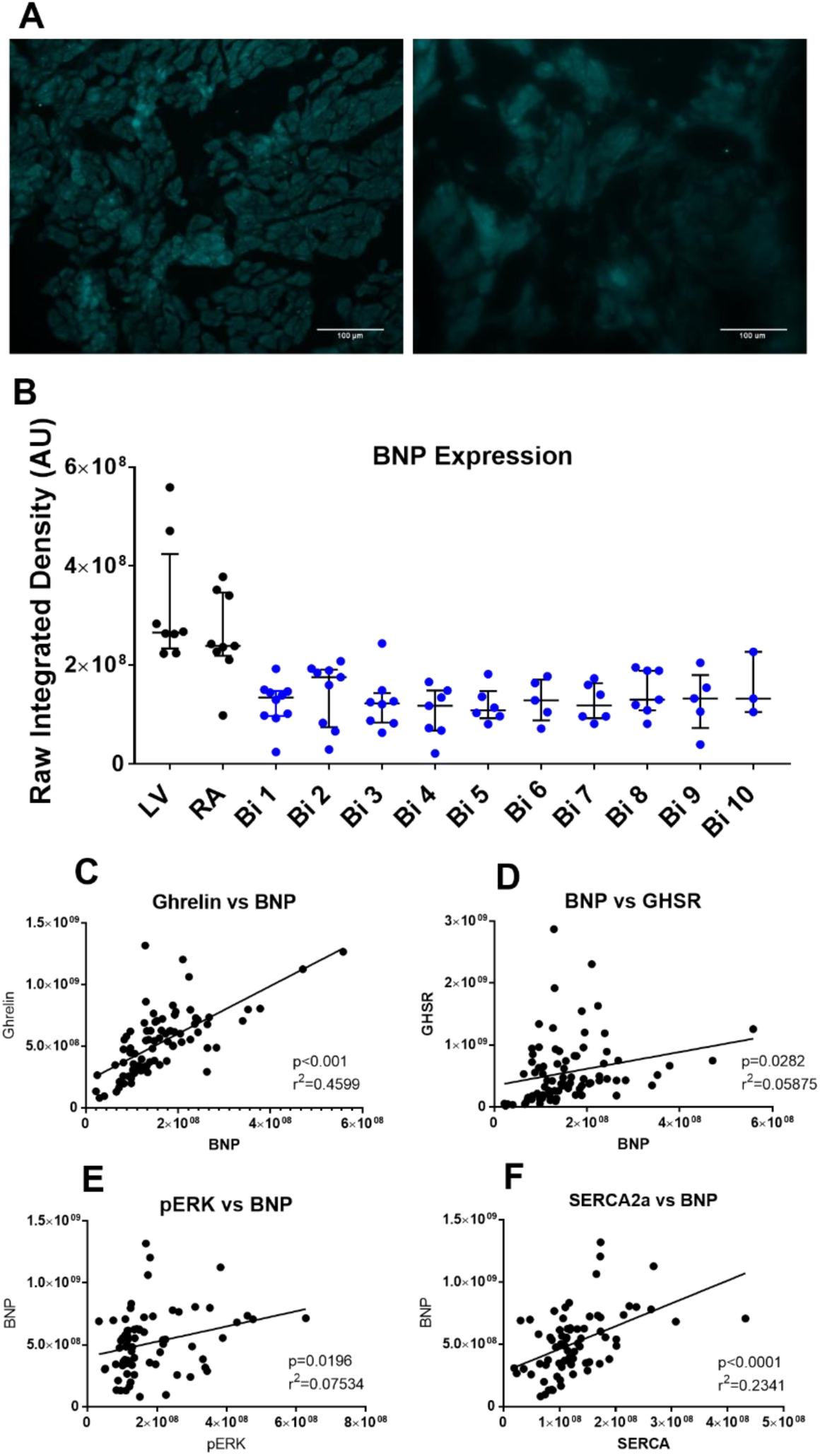
BNP expression in patients pre- and post-cardiac transplant. A) Representative fluorescent images of BNP are shown from explanted heart tissue (left) and grafted heart biopsies (right) taken from the same patient. B) Quantified fluorescence intensity of BNP is shown in explanted heart tissue (LV and RA, black dots) and grafted heart biopsies (Bi 1-10, coloured dots). C-F) Positive correlation is shown in the entire cardiac transplant cohort for ghrelin vs BNP, BNP vs GHSR, pERK1/2 vs BNP, and SERCA2a vs BNP. Each dot represents one transplant patient sample. LV: Left Ventricle; RA: Right atrium; Bi 1 – Bi 10: Biopsy 1 – Biopsy 10.

### 3.4 Cardiac fibrosis

Cardiac fibrosis was determined in all patient samples using Masson’s trichrome stain which measured the presence of collagen I and III (in blue) and compared that to the non-fibrotic tissue (in red). Quantification of fibrosis is illustrated in Figure 4 A, and revealed a high degree of variability both between and within patients from one time point to another (Table 4). Representative images showing the high degree of variability between patients taken from a single time-point are shown in Figure 4 B where significant fibrosis was seen in one patient with large amounts of collagen I and II (blue) and minimal fibrosis was seen in another patient at the same time-point To determine whether GHSR tissue levels were contributed to by fibrosis, cardiac tissue was examined for colocalization between collagen I and GHSR by fluorescence microscopy (Figure 5 A and B). These analyses showed no colocalization between collagen I and GHSR in the human tissue samples (Figure 5C and D).

**Figure 4.**
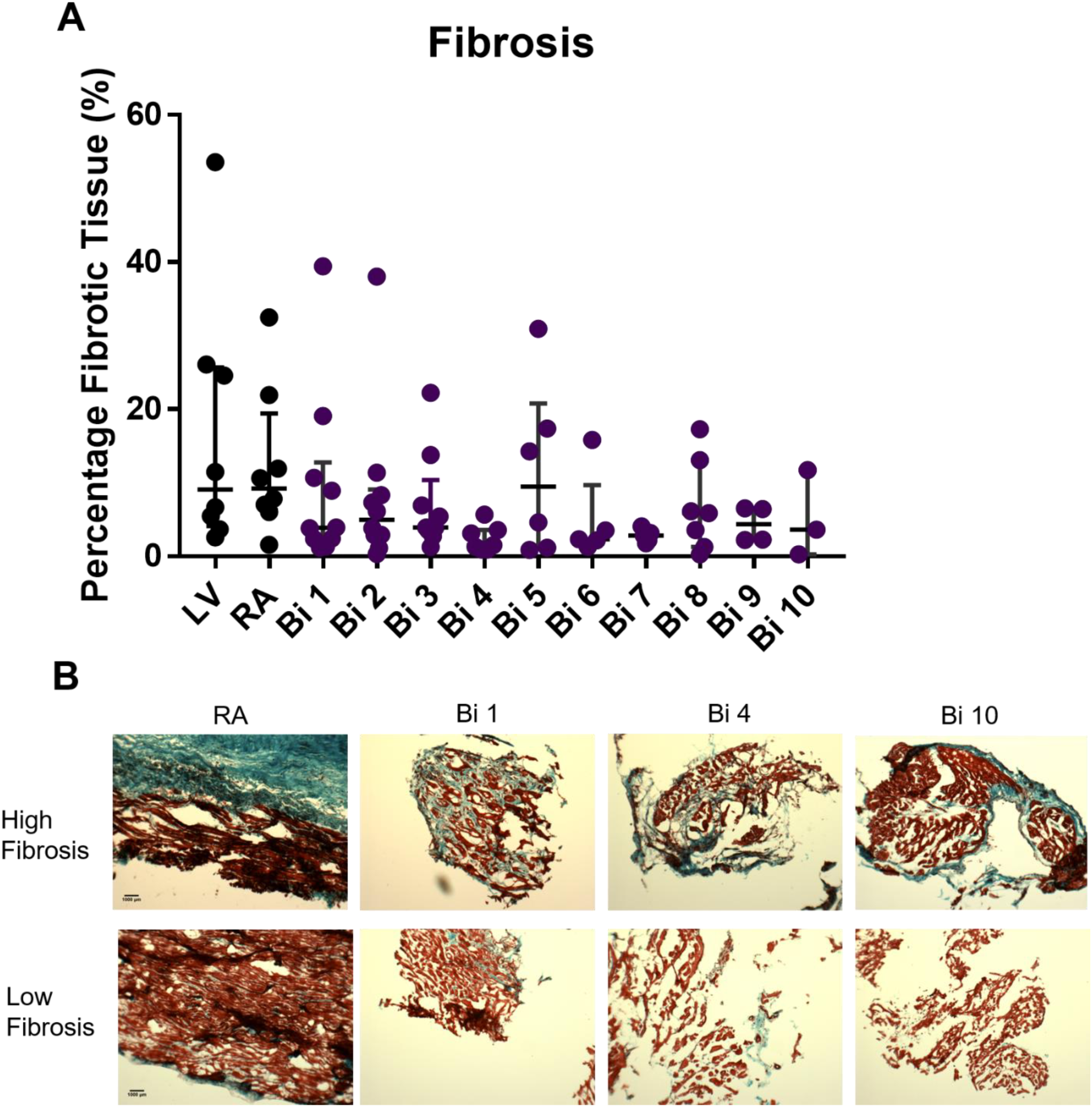
Cardiac fibrosis in patients pre- and post-cardiac transplant. A) Quantified fibrotic data are shown for explanted heart tissue (LV and RA, black dots) and grafted heart biopsies (Bi 1-10, coloured dots). B) Representative images of the fibrotic variability between patients showing high levels of fibrosis (top) and low levels of fibrosis (bottom) where blue is fibrotic tissue (collagen I and III) and red is healthy cardiac tissue. Images of the RA, Bi 1, Bi 4, Bi 10 showing different patients at same time point pre- and post-cardiac transplant. LV: Left Ventricle; RA: Right atrium; Bi 1 – Bi 10: Biopsy 1 – Biopsy 10.

**Figure 5.**
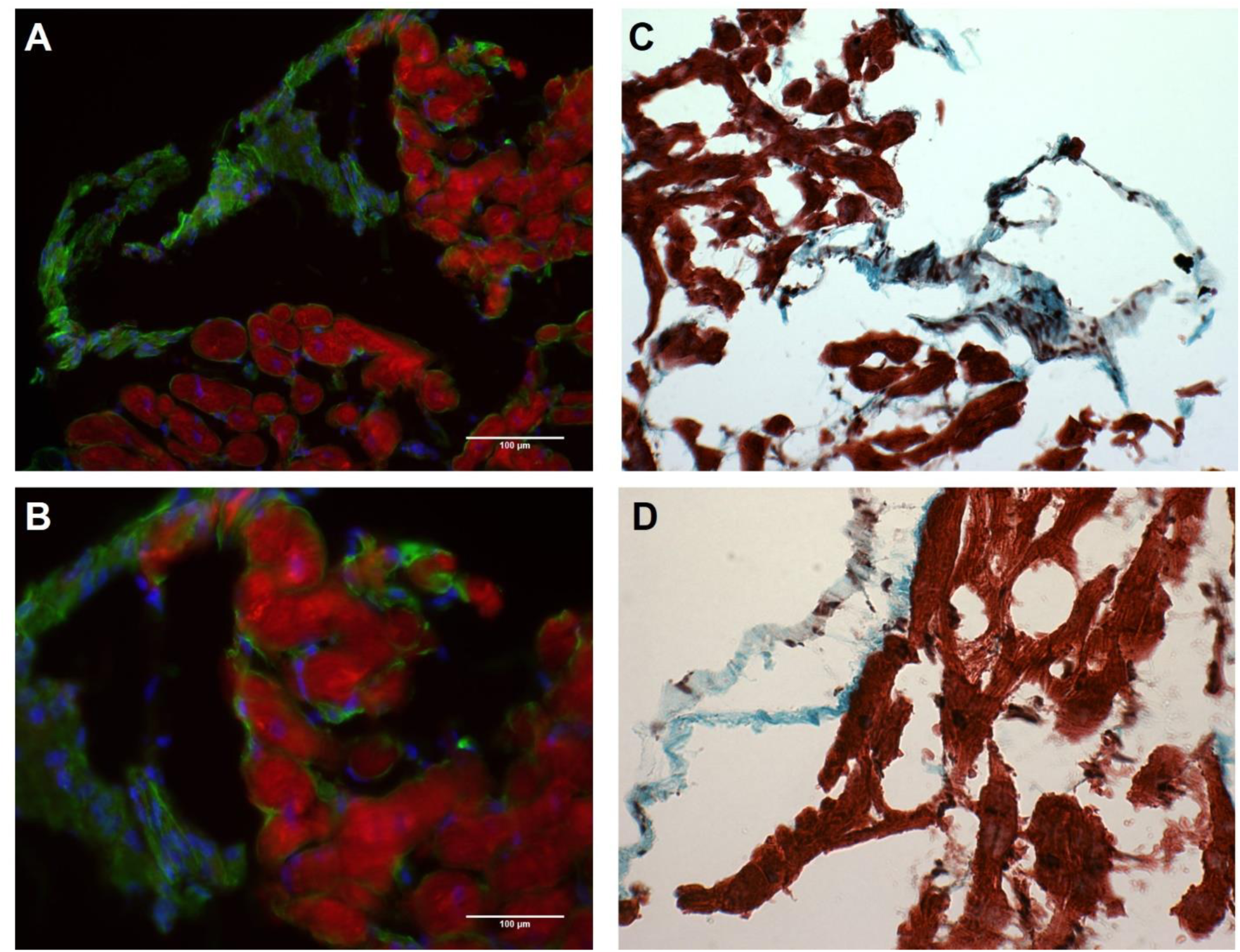
Fibrosis and GHSR in human myocardial tissue. Representative images from grafted heart biopsy 1 showing the same patient sample in all images. A and B show no colocalization of GHSR (red) and collagen I (green) with DAPI showing nuclei (blue). C and D show Masson’s trichrome staining of same patient sample as in A and B where blue is fibrotic tissue (collagen I and III) and red is healthy tissue. A and C show 10X magnification where B and D show 20X magnification.

**Table 4.** Fibrosis Percentages of explanted hearts and healthy implanted heart biopsies. (RA and LV of explanted heart; Biopsy 1-10 of grafted heart). All values listed in percentages (%).

|  | LV | RA | Bi 1 | Bi 2 | Bi 3 | Bi 4 | Bi 5 | Bi 6 | Bi 7 | Bi 8 | Bi 9 | Bi 10 |
| --- | --- | --- | --- | --- | --- | --- | --- | --- | --- | --- | --- | --- |
| Patient 1 | 26.1 | 10.6 | 1.3 | 2.9 | 5.4 | 1.5 | 15.8 | 2.2 | 1.3 | - | - | - |
| Patient 2 | 24.6 | 6.0 | 19.1 | 3.8 | 3.8 | 5.7 | - | - | - | - | - | - |
| Patient 3 | - | 7.0 | 3.9 | 11.4 | 22.2 | 0.9 | - | - | - | 3.6 | 2.2 | - |
| Patient 4 | 2.6 | 32.5 | 1.2 | 0.3 | - | 3.9 | 4.6 | - | - | 13.1 | - | - |
| Patient 5 | 53.6 | - | 2.4 | 7.3 | 3.9 | 1.3 | - | - | - | - | - | - |
| Patient 6 | 3.7 | 7.8 | 2.4 | 38.0 | 1.3 | - | - | - | 4.1 | 5.9 | 2.3 | 11.7 |
| Patient 7 | 5.5 | - | 39.4 | 2.9 | 2.7 | 3.1 | 17.4 | 3.5 | 1.8 | 6.1 | 6.4 | 0.3 |
| Patient 8 | 7.6 | 1.6 | 10.7 | 1.1 | 6.9 | 3.6 | 0.9 | 1.3 | 2.8 | 0.3 | 6.6 | 3.6 |
| Patient 9 | - | 21.9 | 8.9 | 6.1 | 3.5 | - | 1.2 | 2.3 | - | 17.3 | - | - |
| Patient 10 | 11.5 | 11.9 | 3.8 | 8.3 | 13.8 | 0.9 | 30.9 | 2.2 | 3.2 | - | - | - |

## 4. Discussion

This study is the first comprehensive and longitudinal study to examine tissue GHSR, ghrelin and subsequent downstream biochemical signaling molecules involved in cardiomyocyte contractility and growth, over time, in the same patients with two different hearts. Our findings have demonstrated a relationship between cardiac function, as measured by LVEF, and the expression of both ghrelin and GHSR, suggesting that the cardiac ghrelin-GHSR axis may represent a new cardiac-specific biomarker in HF. Our findings further showed an association between the biochemical signaling molecules and their relationship to LVEF. This study is the first to examine the relationship between GHSR, ghrelin, and biochemical signaling molecules in human cardiac tissues and their relationship to LV function, therefore providing substantial new findings related to this myocardial ghrelin/GHSR axis.

Currently the “gold standard” for cardiac biomarkers of heart failure are BNP and NT-proBNP, which are markers of myocardial stress. Historically, past studies using BNP have focused on characterizing HF with reduced EF as associated with and directly linked to elevated circulating BNP levels^12,16^. More recent studies have also attempted to understand the mechanisms leading to HF in the setting of preserved EF. To explore this and other etiologies of HF, a variety of biomarkers including extracellular matrix proteins (matrix metalloproteinase, galectin-3); oxidative stress proteins (8-hydroxy-2,-deoxyguanosine, neopterin); and inflammatory molecules (c-reactive protein, pentraxin 3)^17^ have been identified and their association with the presence and degree of heart failure investigated. Further, circulating biomarkers of HF continue to be discovered through proteomic and gene sequencing of blood serum analyses^18–20^. Many novel serum biomarkers are currently being identified^21,22^, and ghrelin/ GHSR axis may provide a complementary cardiac-specific indicator of LV function in HF. Certainly, biomarkers that are intimately involved in the different pathological processes in HF would help to both enhance early diagnosis and might have the potential to optimize targeted therapies^17,23^. We propose that the ghrelin-GHSR axis represents the first cardiac-specific indicator of HF.

A prior study has shown that GHSR is elevated in chronic heart failure in humans^9^. In contrast to our findings, where both GHSR and ghrelin were elevated, that study also reported a decrease in ghrelin expression in chronic heart failure. The difference in the degree of ghrelin expression between these two studies could reflect the variability in the severity and/or the stage of progression of HF between these two studies. This previously referenced study was also limited by assessment at a single time point with samples only of end stage heart failure^9^. In contrast, our study assessed levels of GHSR and ghrelin at end stage HF and in serial biopsies taken up to a year after transplant surgery, and furthermore, demonstrated significant relationships between LVEF and tissue ghrelin and GHSR levels in two hearts from the same patient. Now that we have established that higher levels of GHSR and ghrelin associate with LVEF in the very low range, these results set the stage for examining changes in cardiac GHSR and ghrelin in patients with a range of LVEF values that reflect the evolution of heart failure.

Circulating ghrelin has also previously been suggested to be a potential single biomarker of HF, and from a prognostic perspective, a level of 85 pmol/l or higher was predictive of increased survival ^24^. However, circulating ghrelin levels are also elevated in the fasting state, and are decreased in both the elderly population and with patients who have a higher BMI^25^, therefore skewing this prognostic cut-off level. In contrast, our data in cardiac tissue samples provide the first direct measure of the ghrelin-GHSR axis being elevated in the human myocardium, thus further strengthening the case for its use as a novel cardiac-specific biomarker of HF within tissue.

As mentioned above, circulating BNP and NT-pro BNP are the currently used clinical biomarkers of HF, as levels rise with decreasing LVEF^26–28^. Our results indicate that levels of BNP in human cardiac tissue trend towards an increase when LVEF is in the very low range, and was more strongly correlated with LVEF than was ghrelin, suggesting that tissue levels of BNP could also be a sensitive biomarker for severe HF. Interestingly, in our study, levels of both GHSR and ghrelin were also associated with LVEF, indicating that the ghrelin-GHSR axis may also be a good cardiac biomarker of LV function. As discussed below, ghrelin showed stronger correlations with biochemical signalling molecules, indicating that tissue ghrelin is more closely associated than is BNP with the biochemical mechanisms that underlie the development of HF.

To better understand the potential mechanisms underlying HF, we examined the downstream signaling pathways that link the ghrelin-GHSR axis to cardiomyocyte contractility and growth. The significant elevation of pERK1/2 levels we observed in end-stage HF is consistent with the reported elevations in pERK1/2 in cardiac tissue in mouse and rat models of HF^29^. Signalling through pERK1/2 is associated with pressure overload-induced myocardial hypertrophy, and thus indicates maladaptive alterations under conditions of persistent myocardial stress. Interestingly, *ghrelin* and *GHSR* gene variants are also associated with LV hypertrophy^30,31^. Therefore, our data suggest a new upregulated ghrelin-GHSR-pERK1/2 pathway that may mediate HF in humans through myocardial hypertrophy.

The strong positive correlation between ghrelin and SERCA2A is in accordance with the role of ghrelin in the improvement of Ca^2+^ dynamics in cardiomyocytes isolated from rodents with ischemia-reperfusion myocardial injury^32^. In this study, activation of GHSR by either ghrelin or hexarelin increased SERCA2a expression and activity through increased phosphorylation of its regulatory binding protein, phospholamban, thus replenishing Ca^2+^ stores in the sarcoplasmic reticulum. However, our results indicating that SERCA2a levels are actually elevated in end-stage HF are in sharp contrast to the literature documenting a decrease in SERCA2a expression and activity in the failing human myocardium^33^. In contrast to the literature, we compared SERCA2a expression between end-stage heart disease and engrafted hearts, and not to healthy controls. The relative decrease in SERCA2a expression in biopsies taken at earlier time points may reflect a subclinical immune response, as a recent study has suggested that decreases in tissue SERCA2a correlate with graft rejection^34^.

Cardiac fibrosis due to a deposition of extracellular matrix proteins including fibroblasts and collagen, occurs in all etiologies of heart disease and heart failure and is a marker of increased heart failure severity^35,36^. Our results indicate an increased collagen I and III deposition in the diseased hearts although there was significant variability both between and within patients. The variability could be a consequence of sampling location; a high presence of fibrosis in the apparently healthy implanted hearts likely indicates a considerable degree of geographic heterogeneity within any given patient’s heart. Traditionally biopsies are acquired from the RV, as what was done here, although a recent study found LV biopsies to be not only possible with low risk via radial access, but preferable for determination of heart function through immunohistochemistry and molecular analyses^37^. Since there was such a large degree of variability in the amount of fibrotic tissue in both pre-transplant hearts and grafted heart tissue, there was a possibility that GHSR expression originated within the fibrotic tissue, thus potentially skewing our results. However, we have shown that GHSR was found only in non-fibrotic tissue; therefore, the variability in expression likely lies in HF type and severity, and not in sampling location. Since the diseased heart tends to have larger amounts of collagen deposition, the positive fluorescent signal is only originating from a limited, non-fibrotic component of the whole tissue section. Therefore, the measurement of GHSR which we used was affected by the extent of collagen deposition within any given sample, with samples with higher degrees of fibrosis having an artificially depressed measurement of “myocardial” levels.

Overall, we have identified the ghrelin-GHSR axis as a cardiac-specific biomarker of cardiac dysfunction in human heart failure. This biomarker was demonstrated to have a stronger sensitivity to the downstream signaling molecules linked to cardiomyocyte contractility and hypertrophy when compared to BNP, the “gold standard” clinically used biomarker. Cardiac fibrosis was highly variable both within and between patients and GHSR was not expressed in fibrotic tissue. We are currently examining the expression and relationship of GHSR-ghrelin signaling that contribute to defective cardiomyocyte programming in other types of heart disease in humans. This work will help to identify GHSR-ghrelin as a cardiac-specific biomarker that can be used to determine the progression of HF at earlier stages.

## Acknowledgements

We like to thank Dr. Peter Pflugfelder, Ms. Anna McDonald and Ms. Stephanie Fox for assistance in the collection of endomyocardial biopsies, and the Pathology Core Lab at the London Health sciences center for their help in fibrosis staining of all tissue samples.

## References

1. Yanagi S, Sato T, Kangawa K, Nakazato M. The Homeostatic Force of Ghrelin. Cell Metab. 2018;27(4):786–804. doi:10.1016/j.cmet.2018.02.008

2. Kishimoto I, Tokudome T, Hosoda H, Miyazato M, Kangawa K. Ghrelin and cardiovascular diseases. Int J Cardiol. 2012;59(1):8–13. doi:10.1016/j.jjcc.2011.11.002

3. Sun Q, Ma Y, Zhang L, Zhao Y-F, Zang W-J, Chen C. Effects of GH Secretagogues on Contractility and Ca2+ Homeostasis of Isolated Adult Rat Ventricular Myocytes. http://dx.doi.org/101210/en2009-1432. 2011. doi:10.1210/EN.2009-1432

4. Douglas GAF, McGirr R, Charlton CL, et al. Characterization of a far-red analog of ghrelin for imaging GHS-R in P19-derived cardiomyocytes. Regul Pept. 2014;54:81–88. doi:10.1016/j.peptides.2014.01.011

5. Baldanzi G, Filigheddu N, Cutrupi S, et al. Ghrelin and des-acyl ghrelin inhibit cell death in cardiomyocytes and endothelial cells through ERK1/2 and PI 3-kinase/AKT. J Cell Biol. 2002;159(6):1029–1037. doi:10.1083/jcb.200207165

6. Sullivan R, McGirr R, Hu S, et al. Changes in the Cardiac GHSR1a-Ghrelin System Correlate With Myocardial Dysfunction in Diabetic Cardiomyopathy in Mice. J Endocr Soc. 2018;2(2):178–189. doi:10.1210/js.2017-00433

7. Yuan M-J, Wang T, Kong B, Wang X, Huang C-X, Wang D. GHSR-1a is a novel pro-angiogenic and anti-remodeling target in rats after myocardial infarction. Eur J Pharmacol. 2016;788:218–225. doi:10.1016/J.EJPHAR.2016.06.032

8. Sullivan R, McGirr R, Hu S, et al. Changes in the Cardiac GHSR1a-Ghrelin System Correlate With Myocardial Dysfunction in Diabetic Cardiomyopathy in Mice. J Endocr Soc. 2018;2(2):178–189. doi:10.1210/js.2017-00433

9. Beiras-Fernandez A, Kreth S, Weis F, et al. Altered myocardial expression of ghrelin and its receptor (GHSR-1a) in patients with severe heart failure. Peptides. 2010;31(12):2222–2228. doi:10.1016/j.peptides.2010.08.019

10. Gaggin HK, Januzzi JL. Biomarkers and diagnostics in heart failure. Mol Basis Dis. 2013;1832:2442–2442. doi:10.1016/j.bbadis.2012.12.014

11. Van Kimmenade RRJ, Januzzi JL. Emerging biomarkers in heart failure. Clin Chem. 2012;58(1):127–138. doi:10.1373/clinchem.2011.165720

12. Braunwald E. Biomarkers in Heart Failure. N Engl J Med. 2008;358(20):2148–2159. http://www.nejm.org.proxy1.lib.uwo.ca/doi/pdf/10.1056/NEJMra0800239. Accessed June 22, 2017.

13. McGirr R, McFarland MS, McTavish J, Luyt LG, Dhanvantari S. Design and characterization of a fluorescent ghrelin analog for imaging the growth hormone secretagogue receptor 1a. Regul Pept. 2011;172(1-3):69–76. doi:10.1016/j.regpep.2011.08.011

14. Guizzetti L, McGirr R, Dhanvantari S. Two dipolar α-helices within hormone-encoding regions of proglucagon are sorting signals to the regulated secretory pathway. J Biol Chem. 2014;289:14968–14980. doi:10.1074/jbc.M114.563684

15. Kennedy DJ, Vetteth S, Periyasamy SM, et al. Central role for the cardiotonic steroid marinobufagenin in the pathogenesis of experimental uremic cardiomyopathy. Hypertension. 2006;47(3):488–495. doi:10.1161/01.HYP.0000202594.82271.92

16. Friões F, Lourenço P, Laszczynska O, et al. Prognostic value of sST2 added to BNP in acute heart failure with preserved or reduced ejection fraction. Clin Res Cardiol. 2015;104(6):491–499. doi:10.1007/s00392-015-0811-x

17. Takeishi Y. Biomarkers in Heart Failure. Int Hear J. 2014;55(6):474–481. https://www.jstage.jst.go.jp/article/ihj/55/6/55_14-267/_pdf. Accessed April 6, 2018.

18. Senthong V, Kirsop JL, Tang WHW. Clinical Phenotyping of Heart Failure with Biomarkers: Current and Future Perspectives. Curr Hear Fail Rep. 2017;14(2):106–116. doi:10.1007/s11897-017-0321-4

19. Meder B, Haas J, Sedaghat-Hamedani F, et al. Epigenome-Wide Association Study Identifies Cardiac Gene Patterning and a Novel Class of Biomarkers for Heart Failure. Circulation. 2017;136(16):1528–1544. doi:10.1161/CIRCULATIONAHA.117.027355

20. Xuan L, Sun L, Zhang Y, et al. Circulating long non-coding RNAs NRON and MHRT as novel predictive biomarkers of heart failure. J Cell Mol Med. 2017;21(9):1803–1814. doi:10.1111/jcmm.13101

21. Stenemo M, Nowak C, Byberg L, et al. Circulating proteins as predictors of incident heart failure in the elderly. Eur J Hear Fail. 2018;20:55–62. doi:10.1002/ejhf.980

22. Van Boven N, Kardys I, van Vark LC, et al. Serially measured circulating microRNAs and adverse clinical outcomes in patients with acute heart failure. Eur J Hear Fail. 2018;20:89–96. doi:10.1002/ejhf.950

23. Bishu K, Redfield MM. Acute Heart Failure with Preserved Ejection Fraction: Unique Patient Characteristics and Targets for Therapy. Curr Hear Fail Rep. 2013;10(3):190–197. doi:10.1007/s11897-013-0149-5

24. Chen Y, Ji X, Zhang A, Lv J, Zhang J, Zhao C. Prognostic value of plasma ghrelin in predicting the outcome of patients with chronic heart failure. J Inst Mex Seg Soc. 2014;45(3):263–269. doi:10.1016/j.arcmed.2014.01.004

25. Rigamonti AE, Pincelli AI, Viarengo R, et al. Plasma ghrelin concentrations in elderly subjects: comparison with anorexic and obese patients. J Endocrinol. 2002;175:1–1. http://joe.endocrinology-journals.org/content/175/1/R1.full.pdf. Accessed April 23, 2018.

26. Karakilic E, Kepez A, Abali G, Coskun F, Kunt M, Tokgozoglu L. The relationship between B-type natriuretic peptide levels and echocardiographic parameters in patients with heart failure admitted to the emergency department. Anadolu Kardiyol Dergisi/The Anatol J Cardiol. 2010;10(2):143–149. doi:10.5152/akd.2010.040

27. Jiang Yanxia MC, Yanxia J, Xingjun C, Xintao T, Lei S. The Correlation between Left Ventricular Ejection Fraction and Peripheral Blood MCP-1 NT-Pro BNP in Patients with Acute Coronary Syndrome. Intern Med Open Access. 2014;04(01):1–4. doi:10.4172/2165-8048.1000139

28. Belagavi AC, Rao M, Pillai AY, Srihari US. Correlation between NT proBNP and left ventricular ejection fraction in elderly patients presenting to emergency department with dyspnoea. Indian Heart J. 2012;64(3):302–304. doi:10.1016/S0019-4832(12)60091-1

29. Bueno OF, Molkentin JD. Involvement of extracellular signal-regulated kinases 1/2 in cardiac hypertrophy and cell death. Circ Res. 2002;91(9):776–781. doi:10.1161/01.RES.0000038488.38975.1A

30. Baessler A, Kwitek AE, Fischer M, et al. Association of the ghrelin receptor gene region with left ventricular hypertrophy in the general population: Results of the MONICA/KORA Augsburg echocardiographic substudy. Hypertension. 2006;47(5):920–927. doi:10.1161/01.HYP.0000215180.32274.c8

31. Ukkola O, Pääkkö T, Kesäniemi YA. Ghrelin and its promoter variant associated with cardiac hypertrophy. J Hum Hypertens. 2012;26(7):452–457. doi:10.1038/jhh.2011.51

32. Ma Y, Zhang L, Edwards JN, et al. Growth Hormone Secretagogues Protect Mouse Cardiomyocytes from in vitro Ischemia/Reperfusion Injury through Regulation of Intracellular Calcium. Lewin A, ed. PLoS one. 2012;7(4):e35265. doi:10.1371/journal.pone.0035265

33. Gorski PA, Ceholski DK, Hajjar RJ. Altered myocardial calcium cycling and energetics in heart failure - A rational approach for disease treatment. Cell Metab. 2015;21(2):183–194. doi:10.1016/j.cmet.2015.01.005

34. Tarazón E, Ortega A, Gil-Cayuela C, et al. SERCA2a: A potential non-invasive biomarker of cardiac allograft rejection. J Hear Lung Transplant. 2017;36(12):1322–1328. doi:10.1016/j.healun.2017.07.003

35. Majumder R, Nayak AR, Pandit R. Nonequilibrium Arrhythmic States and Transitions in a Mathematical Model for Diffuse Fibrosis in Human Cardiac Tissue. Chen X, ed. PLoS one. 2012;7(10):e45040. doi:10.1371/journal.pone.0045040

36. Boldt A, Wetzel U, Lauschke J, et al. Fibrosis in left atrial tissue of patients with atrial fibrillation with and without underlying mitral valve disease. BMJ Hear. 2004;90(4):400–405. doi:10.1136/HRT.2003.015347

37. Frey N, Meder B, Katus HA. Left Ventricular Biopsy in the Diagnosis of Myocardial Diseases. Circulation. 2018;137(10):993–995. doi:10.1161/CIRCULATIONAHA.117.030834

